# Diffusion-Based Inference for Heterogeneous Multichannel Fluorescence Imaging

**DOI:** 10.64898/2026.10.07.753438

**Authors:** Ioannis Georvasilis, Marcos Frías Nestares, Carmen Badosa, Uliana Musokhranova, Alfonso Oyarzábal, Cecilia Jimenez-Mallebrera, Jacek Rumiński, Mònica Roldán

## Abstract

Fluorescence microscopy is a central tool in biomedical image analysis, but its practical use is often constrained by incomplete and heterogeneous data. This is especially true in rare disease research, where patient cohorts are small, biological material is limited, and complex study designs frequently yield images with missing, degraded, or low-quality channels. We introduce Reliability-Mask-Conditioned DDPM (RM-DDPM), a conditional diffusion model that formulates multichannel image generation, missing-channel imputation, and degraded-channel restoration as a single reliability-guided inference problem. The model is trained on a curated subset of complete, high-quality multichannel images using a self-supervised masking objective to learn rich cross-channel dependencies from the available channels. At inference time, reliable observations condition the reverse diffusion process, allowing the same trained model to solve multiple reconstruction tasks without task-specific retraining. We evaluate RM-DDPM against task-specific baselines on one public and two in-house fluorescence microscopy datasets, comprising a rare-disease preclinical model study and a clinical patient cohort. Across these settings, RM-DDPM matches the specialized baselines in reconstruction quality, hallucinates fewer false structures, and remains more robust as degradation severity increases, while repeated sampling provides per-pixel uncertainty maps that flag unreliable reconstructions.

## 1. Introduction

Multichannel fluorescence microscopy is a key tool in biological and medical imaging, enabling simultaneous visualization of multiple cellular structures, molecular pathways, and functional processes within a single specimen. By combining complementary fluorescence channels, these acquisitions provide richer biological context, multiplexed biomarker discovery, and greater robustness for downstream and representation-learning workflows compared to single-channel imaging (1–3).

Despite these advantages, fully observed and standardized multichannel datasets remain uncommon, particularly in rare disease research, where biological specimens are often difficult to reproduce, datasets are small, and acquisition protocols evolve over time (4, 5). Earlier samples may therefore contain fewer fluorescence channels than more recent acquisitions, leading to incomplete observations. In addition, channel quality can vary due to photobleaching, low signal-to-noise ratios, optical blur, staining variability, or protocol-specific artifacts. Such variability may occur across samples as well as within the same specimen across different regions or fields of view (5, 6). Figure 1 illustrates representative forms of biological, acquisition, and channel-level heterogeneity observed in one of the two real-world fluorescence microscopy datasets considered in this work.

**Fig. 1.**
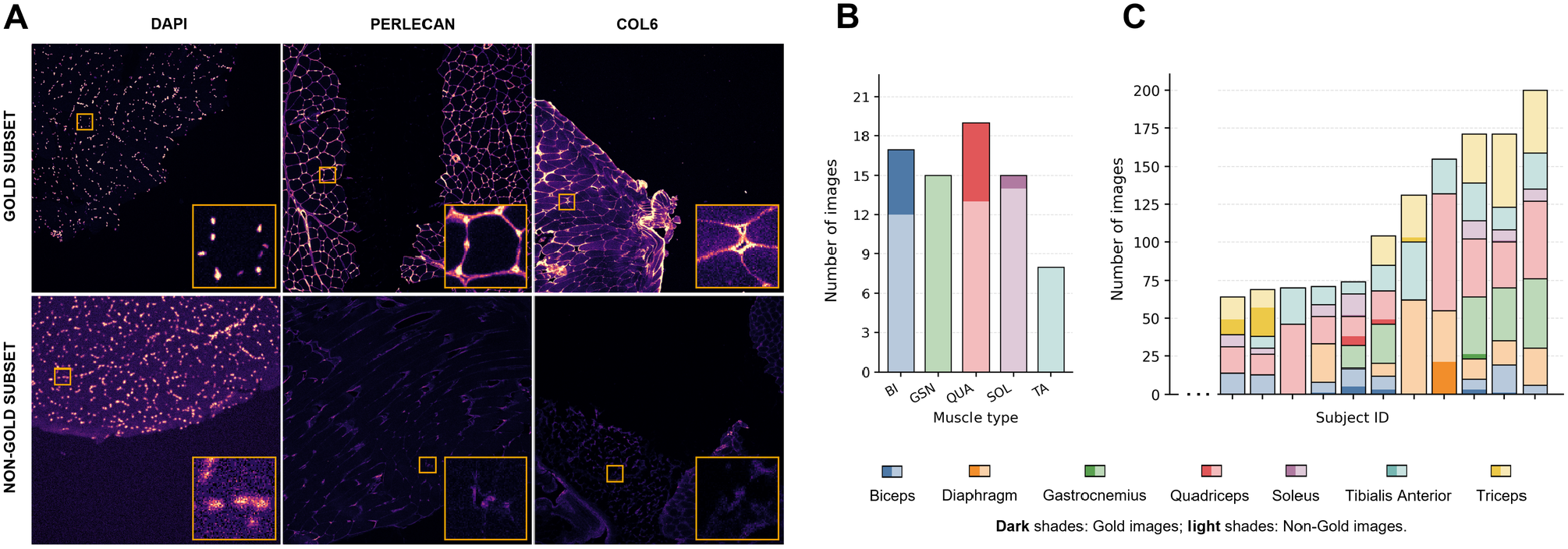
Heterogeneity and gold-standard curation in Col6a1-KI. (A) Representative gold-standard vs. non-gold images across the three channels (DAPI, Perlecan, Collagen VI); insets highlight degradation artifacts. (B) Gold vs. non-gold image counts by muscle type for one subject. (C) Same breakdown across all subjects (dark = gold, light = non-gold).

These limitations directly affect downstream computational analysis. Many image processing pipelines (e.g., segmentation, feature extraction, representation learning) assume fixed channel configurations and reliable signal across modalities (5, 7). Samples with missing or degraded channels are therefore often excluded, reducing dataset size and potentially introducing bias. This issue is particularly severe in rare disease studies, where each sample is uniquely valuable (4). In practice, only a limited subset of samples may be fully observed and consistently high quality across all channels. In this work, we refer to this curated subset as the gold-standard subset, which we adopt as the high-quality reference for modeling complete multichannel observations.

Addressing this setting requires methods that can learn from limited high-quality multichannel data while remaining robust to heterogeneous missingness and degradation at inference time. Existing approaches address related aspects of the problem, including multichannel image generation, missing-channel imputation, and degraded-channel restoration. However, many are designed around predefined source– target channel configurations (8, 9), treat channels independently (10, 11), assume access to a large number of channels with rich cross-channel correlations (12, 13), or rely on specific degradation models (14, 15), limiting their applicability when channel availability and signal reliability vary across samples (16).

To address these limitations, we propose Reliability-Mask-Conditioned Denoising Diffusion Probabilistic Model (RM-DDPM), an approach for heterogeneous multichannel fluorescence microscopy. Rather than treating multichannel image generation, missing-channel imputation, and degraded-channel restoration as independent problems requiring separate models, RM-DDPM formulates them as different instances of the same reliability-guided reconstruction problem. We assume access to, or the ability to extract, a gold-standard subset of complete, high-quality multichannel images, on which the model is trained using a self-supervised masking objective to learn a structured prior over biologically plausible microscopy data.

At inference, the model is conditioned on a pixel-wise reliability mask indicating which observations are trustworthy and which are missing or degraded. This allows a single trained model to handle arbitrary combinations of missing channels and unreliable regions, without requiring separate models for each scenario.

The main contributions of this work are:

- We formulate heterogeneous multichannel fluorescence reconstruction as conditional inference problem over a learned gold-standard image distribution, encompassing multichannel image generation, missing-channel imputation, and degraded-channel restoration.
- We introduce RM-DDPM, a diffusion model that uses a reliability mask to condition image reconstruction, allowing a single trained network to handle arbitrary combinations of missing channels and degraded regions without task-specific retraining, while explicitly grounding the reverse process in reliable observations to mitigate hallucination relative to unconditioned baselines.
- We evaluate the framework across controlled and real-world acquisition settings, showing that a single reliability-conditioned prior remains competitive with task-specific supervised models, hallucinates fewer false structures, and is more robust as degradation severity increases.

## 2. Related Work

### 2.1. Missing-channel Imputation

Numerous methods have been proposed for missing-channel imputation. Most of these methods are closely related to virtual staining and in silico labeling, where computational models predict unobserved fluorescence markers from a subset of acquired images or from alternative imaging modalities. Early approaches used supervised regression and encoder–decoder convolutional neural networks to establish deterministic mappings between the available inputs and target channels (8, 9).

To capture complex cross-channel dependencies, subsequent methods moved beyond simple regression toward architectures explicitly designed for cross-channel modeling. Adversarial, attention-based, and image-to-image translation methods have been used to synthesize realistic fluorescence signals or virtual stains from label-free, partial, or related imaging inputs (17–19). Current strategies integrate masking-based training and reduced-panel learning to improve architectural robustness and mitigate protocol variability (12, 13, 20). These methods are predominantly evaluated on multiplex immunofluorescence datasets, where numerous simultaneously acquired markers provide rich cross-channel information for predicting missing fluorescence channels.

Together, these approaches demonstrate that fluorescence channels can often be inferred from correlated structural or molecular information. However, many imputation and virtual staining methods remain optimized for predefined source–target configurations, reduced marker panels, or settings with a large number of strongly correlated channels. Moreover, most methods assume reliable input channels, limiting their applicability when observations may be simultaneously missing, noisy, blurred, attenuated, or otherwise unreliable.

### 2.2. Degraded-channel Restoration

Beyond missing channels, fluorescence microscopy acquisitions routinely suffer from channel-specific degradation. Artifacts such as photon noise, optical blur, photobleaching, and staining inconsistencies obscure critical cellular structures, directly compromising the robustness of subsequent quantitative analyses.

Classical restoration relied on statistical signal processing and model-based reconstruction. Representative approaches include variance-stabilizing transforms for Poisson noise, non-local means and total variation regularization for denoising, and Richardson–Lucy deconvolution for optical blur (21–24). Deep learning methods subsequently advanced restoration by learning mappings from degraded observations to high-quality targets. When paired training data are available, supervised networks such as CARE (10), RCAN (14), and DFCAN (15) can recover degraded fluorescence signals. To relax the requirement for clean reference images, self-supervised and unsupervised restoration methods exploit internal image structure and noise statistics to estimate the underlying signal (11, 25–29).

Despite these advances, existing restoration methods typically rely on restrictive assumptions about the observation process, including single-channel inputs, fixed sets of fully observed modalities, fixed clean–degraded training pairs, or degradation-specific models tailored to individual artifact types. These assumptions are often violated in heterogeneous multichannel fluorescence datasets, where channel availability and degradation patterns vary across samples, fields of view, and acquisition protocols.

### 2.3. Diffusion Models in Medical Imaging

Diffusion models have emerged as a powerful generative framework, learning complex image distributions via a gradual denoising process (30, 31). Beyond unconditional generation, diffusion models have been extended to image-to-image translation and inverse problems through conditional formulations and guided sampling strategies (32–34). These approaches enable reconstruction from incomplete or degraded observations by enforcing consistency with available measurements during the reverse process.

In biomedical imaging, diffusion models have shown strong performance across modalities such as MRI, CT, ultrasound, and digital pathology, supporting both data generation and reconstruction tasks (35–40). Conditional diffusion frameworks have also been applied to cross-modality synthesis and restoration, demonstrating the potential of probabilistic modeling for handling incomplete observations (41).

In fluorescence microscopy, diffusion-based approaches are only beginning to emerge. Recent work applies diffusion models to unpaired fluorescence microscopy denoising (42), while related approaches explore diffusion-based synthesis and virtual staining (43–45).

Despite these advances, most diffusion-based approaches remain tied to specific observation models, such as denoising an observed fluorescence image, reconstructing from a known measurement operator, or translating autofluorescence inputs into predefined stained targets. Consequently, they do not naturally accommodate heterogeneous multi-channel fluorescence datasets, where channel-wise and spatial reliability patterns can vary across samples and fields of view. This motivates our formulation of reconstruction as conditional inference over a learned gold-standard distribution.

## 3. Methods

In this section, we formulate multichannel fluorescence reconstruction under heterogeneous missingness and degradation. We begin by defining the curated gold-standard subset used to learn the multichannel image prior, then introduce the RM-DDPM framework and its conditional inference procedure.

### 3.1. Gold-Standard Subset Definition

We consider a multichannel fluorescence microscopy dataset 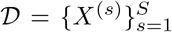, where *S* denotes the total number of specimens and each specimen 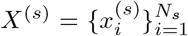 contains *N*_*s*_ spatially aligned multichannel images 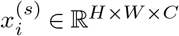, with *C* denoting the number of fluorescence channels. In addition, channel availability and signal quality may vary both across and within specimens. Common sources of degradation include measurement noise, optical blur, intensity attenuation due to staining variability, axial drift, and photobleaching. The complete collection of individual images across all specimens is therefore

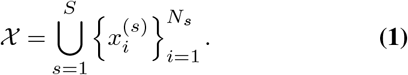

Our goal is to learn a generative prior over clean, fully observed multichannel images. We therefore define the gold-standard subset as

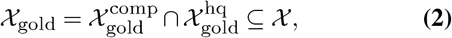

where 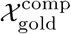 denotes the set of images with all required fluorescence channels available (complete), and 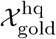 denotes the set of images that pass image-based quality control and are therefore considered high quality.

Selection into *X*_gold_ is performed per image rather than per specimen, since channel availability and quality can vary from image to image within the same specimen. Consequently, a given specimen may contribute a subset of its images, while the remaining images are excluded.

Channel completeness is determined from acquisition metadata, whereas image quality is assessed directly from the individual fluorescence channel images in the original heterogeneous dataset. In general, the gold-standard subset may be identified through automated quality control, expert curation, or a combination of both. In this work, we adopt an automated image-based quality-control procedure based on simple, interpretable image quality metrics established for microscopy (46, 47).

For each channel *c*, we define a dataset-specific multi-objective quality function Φ_*c*_(*x*_*c*_) composed of image quality metrics (IQMs). A channel is considered reliable if it satisfies the corresponding quality criterion

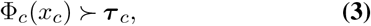

where ***τ***_*c*_ denotes the channel-specific quality thresholds. The dataset-specific IQMs and threshold values are reported in Appendix A.

**Fig. 2.**
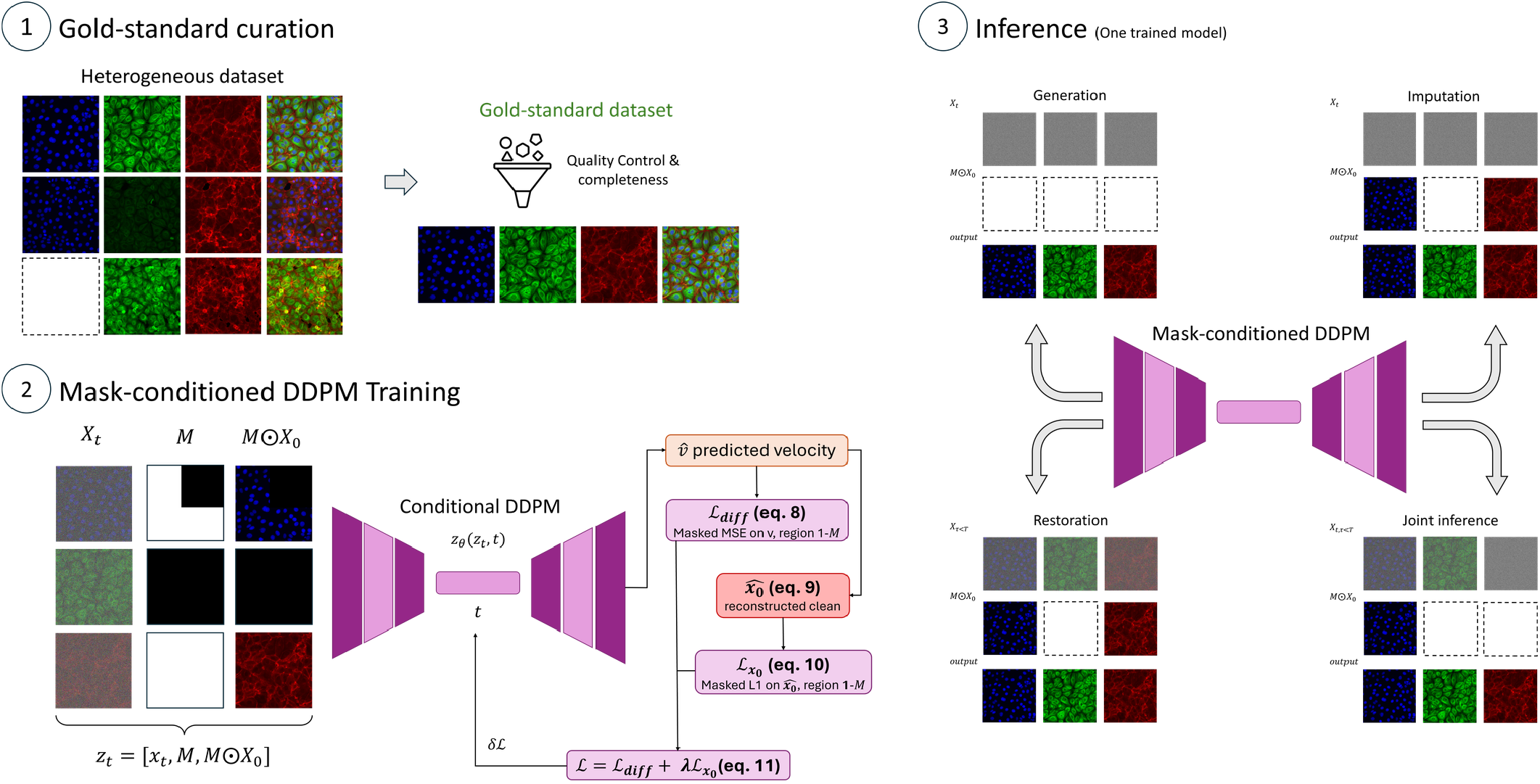
Method overview. (1) *Gold-standard curation*: a quality-control and completeness step selects the gold-standard subset from the heterogeneous dataset. (2) *Mask-conditioned DDPM training*: the network is trained on masked inputs *z*_*t*_ = [*x*_*t*_, *M, M ⊙ x*_0_] from gold-standard images, optimizing a masked diffusion loss and an auxiliary *x*_0_-reconstruction loss over the unreliable region. (3) *Inference*: the same trained model performs generation, imputation, restoration, and joint inference by changing only the reliability mask and context.

This selection yields a subset of high-quality, complete multichannel fluorescence images with zero missingness and reduced degradations.

### 3.2. Reliability-Mask-Conditioned DDPM

We propose RM-DDPM to learn the distribution of multichannel images in the gold-standard subset. Let *x*_0_ ~*pX*_gold_denote a complete, high-quality multichannel image sampled from the gold-standard subset. The forward diffusion process gradually corrupts *x*_0_ with Gaussian noise according to

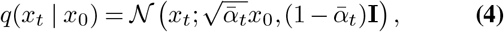

where 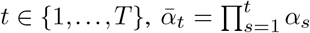, and 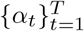 defines the noise schedule.

RM-DDPM is trained using the velocity parameterization (48). Given Gaussian noise *ϵ*~ *N*(0, **I**), the noised sample is written as

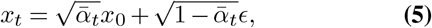

and the velocity target is defined as

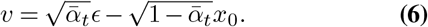

A neural network *v*_*θ*_ is trained to predict this velocity target. To reconstruct partial and heterogeneous observations, RMDDPM is conditioned on a binary reliability mask *M* ∈ {0, 1}^*H×W×C*^, where *M* = **1** denotes reliable context, whereas *M* = **0** denotes components that the model must infer.

During training, reliability masks are synthetically generated by masking gold-standard images. The network input is the channel-wise concatenation of the noised sample, the reliability mask, and the masked gold-standard image, which serves as the reliable context and is replaced at inference by the actually observed reliable channels (Section 3.4):

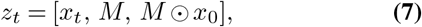

where ⊙ denotes element-wise multiplication.

The diffusion loss is evaluated only over the unreliable components:

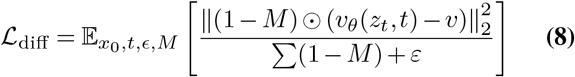

where *ε* is a small constant used to avoid division by zero. From the predicted velocity *v*_*θ*_, we obtain an estimate of the clean sample 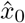

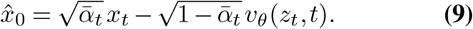

To directly supervise the unreliable components, we incorporate an auxiliary reconstruction term evaluated only over the target region (1 − *M*):

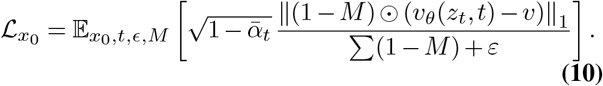

The final training objective is

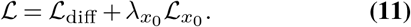

This combined objective encourages the model to reconstruct unreliable regions while preserving the reliable context.

### 3.3 Masking Strategy

During training, the reliability masks *M* are sampled uniformly from a predefined mixture of reconstruction scenarios to expose the model to diverse reliability patterns. This mixture comprises channel-wise masking, spatial masking, and joint channel-spatial masking. Specifically, channel-wise masks simulate missing fluorescence modalities, spatial masks simulate localized artifact corruption, and mixed masks capture heterogeneous degradation patterns. To enforce learning across all tasks, the fully unobserved configuration (*M* = **0**) is included to retain unconditional generation capabilities, whereas the fully observed configuration (*M* = **1**) is explicitly excluded as a trivial identity mapping. Detailed definitions and configurations for each masking case are provided in Appendix B

### 3.4 Conditional Inference via Masked Diffusion

Having learned a prior over complete, high-quality multichannel images, we now describe how the same trained model performs inference under heterogeneous reliability patterns. At inference time, the trained model performs conditional sampling given whatever reliable observations are available, guided by the prior learned over the gold-standard distribution.

Let *x*_ctx_ ∈ ℝ^*H×W×C*^ denote the available reliable context, with missing or unreliable channels zero-filled so that *x*_ctx_ always retains *C* channels. Let *M* ∈ {0, 1}^*H×W×C*^ be the corresponding reliability mask, as defined in Section 3.2, with *M* = 1 components preserved throughout sampling.

At each reverse diffusion step, the network is conditioned in the same way as during training, receiving as input

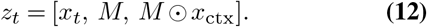

The model predicts the velocity *v*_*θ*_(*z*_*t*_, *t*), which parameterizes the reverse transition and yields an intermediate estimate 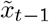:

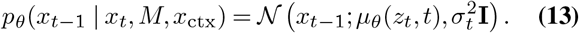

To enforce consistency with the reliable context, we replace the reliable components after each reverse step with the corresponding forward-noised version of the reliable observation, as in RePaint (33):

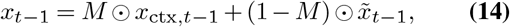

where

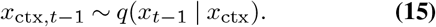

At the final timestep, the reliable components are copied from *x*_ctx_, while the unreliable components are taken from the generated sample:

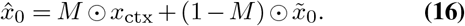

Throughout, the model is referred to as RM-DDPM when sampled with the ancestral DDPM sampler and as RMDDIM when using the deterministic Denoising Diffusion Implicit Models (DDIM) sampling procedure (49).

Different inference regimes are specified by the reliability mask *M*, context image *x*_ctx_, and starting timestep *t*_start_:

- **Multichannel image generation**. When no reliable context is available, we set *M* = **0** and initialize the trajectory from Gaussian noise, *x*_*T*_~ *N* (0, **I**). The model then samples a complete multichannel image from the learned gold-standard prior.
- **Missing-channel imputation**. For a subset of missing target channels *C*_miss_ ⊂ {1,…, *C*}, we set *M*_*c*_ = 0 for all *c* ∈ *C*_miss_ and 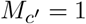 for the remaining available reliable channels *c*′ ∉*C*_miss_. The reverse process is initialized from noise at *t*_start_ = *T*. The model jointly reconstructs the missing channels conditioned on the reliable channels, while the consistency step preserves the reliable context.
- **Degraded-channel restoration**. For a subset of degraded target channels *C*_deg_ ⊂ {1,…, *C}*, we set *M*_*c*_ = 0 for all *c* ∈ *C*_deg_ and 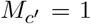 for the remaining reliable channels *c*′ ∉*C*_deg_. Unlike imputation, the reverse trajectory is not started from noise but from the degraded channels partially noised to an intermediate timestep *t*_start_ = *τ < T*, following the partial-noising initialization of SDEdit (50). This preserves their coarse structure while leaving the reverse process free to remove corruption and jointly project the degraded channels toward the gold-standard distribution. Reliable non-target channels are preserved by the consistency step. The choice of *τ* trades measurement fidelity against prior strength: values near *T* favor the learned prior and risk hallucination, as in imputation, while values near 0 retain too much of the degraded signal to remove the corruption. The specific *τ* values used for each dataset and task are reported in Section 4.3.
- **Unconditional restoration**. When no channel is treated as reliable, *M* = **0** as in unconditional generation, but instead of initializing from pure noise, the reverse trajectory is initialized by partially noising the degraded observation to an intermediate timestep *t*_start_ = *τ < T*, as in degraded-channel restoration. Since *M* = **0** everywhere, no channel serves as reliable context for the others and the consistency step has no effect. The partially noised degraded observation is the only sample-specific input, and the reverse process relies entirely on the learned prior to jointly restore all channels.

Thus, each inference regime is specified through the triplet (*M, x*_ctx_, *t*_start_), without task-specific retraining.

Because the reverse process above is stochastic, it can be run multiple times for the same input under independent noise realizations. Averaging *K* such reconstructions (RM-DDPM ×*K* / RM-DDIM× *K*) improves the point estimate, while the pixel-wise variance across the *K* samples yields a per-pixel uncertainty map at no additional training cost, providing a built-in uncertainty estimate over the reconstructed unreliable regions.

## 4. Experiments and Evaluation

We evaluate the proposed framework across two complementary settings. First, we use an external fully observed fluorescence microscopy dataset to quantify reconstruction performance under synthetic missingness and channel degradation, where withheld or degraded channels remain available as paired references. Second, we use heterogeneous in-house rare-disease fluorescence microscopy datasets without paired clean ground truth to assess practical behavior under limited sample availability, variable acquisition quality, and incomplete or degraded channels.

### 4.1. Datasets

All datasets considered in this study contain three fluorescence channels (*C* = 3), although the corresponding markers differ across datasets. Table 1 summarizes their modalities, subset sizes, gold-standard availability, and spatial dimensions. Figure 3 shows a representative example from each dataset, illustrating their individual fluorescence channels together with the composite overlay.

**Table 1.** Dataset summary. Size = number of images (BBBC021v1, Dnm1L-Mito) or mosaics (Col6a1-KI); #Gold = count (%) of gold-standard samples, same unit as Size (tile-level rate for Col6a1-KI reported in Section 4.1). Tasks: Gen = generation, Imp = imputation, Rest = restoration.

| Dataset | Modalities | Size | #Gold (%) | Task |
| --- | --- | --- | --- | --- |
| BBBC021v1 | DAPI, Tubulin, Actin | 240 | 24 (10%) | Gen, Imp, Rest |
| Col6a1-KI | DAPI, Perlecan, Collagen VI | 184 | 23 (12.5%) | Gen, Rest |
| Dnm1L-Mito | DAPI, α-tubulin, TOMM20 | 77 | 73 (95%) | Gen, Imp |

**Fig. 3.**
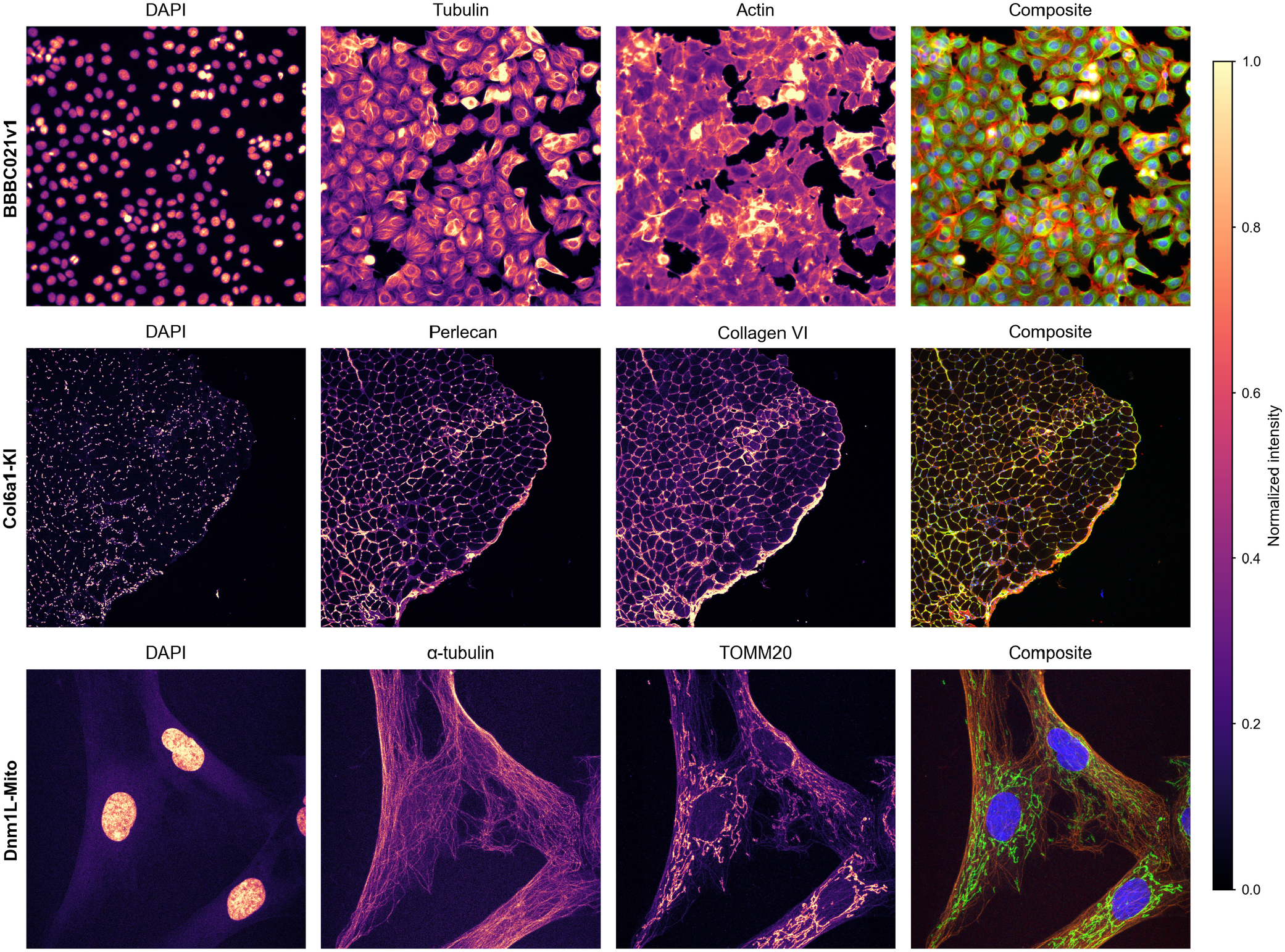
Representative dataset examples. One representative image is shown for each dataset (BBBC021v1, Col6a1-KI, and Dnm1L-Mito; rows), together with the corresponding individual fluorescence channels and their composite overlay (columns).

#### External controlled evaluation dataset

We use the Week10_40111 subset of BBBC021v1 (51, 52) as an external, fully observed benchmark, comprising 240 multichannel images (1024× 1280 px) of MCF-7 breast cancer cells with DAPI, Tubulin, and Actin channels. We deliberately select this three-channel dataset—rather than one with more imaging channels—to emulate the low-channel-count acquisition protocols common in rare-disease studies. Its complete channel availability enables controlled quantitative evaluation via synthetic channel withholding or degradation against paired reference targets. To further emulate the limited curated-data availability of heterogeneous rare-disease studies, we treat only a randomly selected 10% of BBBC021v1 as the available gold-standard subset across all experiments—a conservative, low-curation scenario not intended to match any single real-world dataset’s gold-standard rate.

#### Real-world heterogeneous datasets

Real-world validation is performed on rare-disease fluorescence microscopy datasets characterized by limited gold-standard availability, heterogeneous acquisition quality, and channel-specific degradation.

The Col6a1-KI skeletal muscle dataset (53) acquired by the Neuromuscular Unit at Hospital Sant Joan de Déu, consists of 184 multichannel mosaics containing DAPI, Perlecan, and Collagen VI channels. For Col6a1-KI, quality control was performed at the tile level because individual mosaics may contain both high-quality and degraded regions. The 184 mosaics were decomposed into fixed-size 1024 × 1024 px tiles, and channel-specific quality criteria were applied independently to each tile. This process identified 113 high-quality gold-standard tiles, originating from 23 mosaics, among the 1,877 tiles evaluated. To prevent subject-level leakage, the 23 mosaics contributing gold-standard training tiles are disjoint from those providing the non-gold tiles used for restoration evaluation. Appendix A reports the resulting per-tile quality control outcomes and channel-specific criteria applied to this dataset.

The Dnm1L-Mito, acquired by the Synaptic Metabolism and Personalized Therapies Lab at Hospital Sant Joan de Déu, originates from primary skin fibroblasts of DNM1Lpatients, patients with other mitochondrial disorders, and matched controls, used to characterize mitochondrial network morphology—an established phenotypic biomarker for DNM1L-related mitochondrial fission deficiency (54, 55). It comprises 77 multichannel fluorescence microscopy images containing DAPI, *α*-tubulin, and TOMM20 channels, corresponding to nuclei, cytoskeleton, and the mitochondrial network, respectively. Unlike Col6a1-KI, the Dnm1L-Mito dataset is natively acquired at tile resolution rather than as larger mosaics. All images have a fixed spatial resolution of 1024× 1024 px. Among them, 73 contained complete channel acquisitions and were retained as gold-standard reference samples.

### 4.2. Evaluation Protocol

We evaluate the proposed framework under two complementary regimes: controlled experiments on BBBC021v1, where missing channels, synthetically degraded channels, and their combination can be assessed against paired references, and real-world experiments on the two rare-disease datasets, where no paired clean reference exists and surrogate criteria are required.

#### Compared methods

The same set of diffusion models is compared across all tasks. Baseline DDPM and Baseline DDIM denote plain diffusion baselines, sampled with ancestral DDPM sampling (30) and 50-step DDIM sampling (49), respectively. RM-DDPM and RM-DDIM denote the proposed context-conditioned reliability-masked variants defined in Section 3.2. RM-DDPM ×10 and RM-DDIM ×10 apply the stochastic ensembling described in Section 3.4, averaging 10 independent samples per input to form the final prediction and deriving a pixel-wise uncertainty map from their variance. Task-specific baselines are introduced alongside each task below.

#### Evaluation metrics

Two metric families are used, depending on whether a pixel-aligned reference is available. For imputation and restoration tasks with a paired reference, reconstruction quality is assessed with Structural Similarity Index Measure (SSIM) (56), Learned Perceptual Image Patch Similarity (LPIPS, AlexNet backbone) (57), and Cellpose-SAM-derived matched-object Dice scores (58). For generation, where no pixel-aligned target exists, distributional quality is quantified with the F1 score from Improved Precision and Recall (iF1) (59), Fréchet Inception Distance (FID) (60), and Kernel Inception Distance (KID) (61), all computed in a channel-aware biological embedding space (Appendix C). On the real-world datasets neither family applies directly, since no paired reference exists for any task; the surrogate criteria used there are described in Real-world evaluation below.

#### Controlled evaluation (BBBC021v1)

Four tasks are evaluated under this fully observed, paired-reference setting.

##### Unconditional image generation

Generated samples are compared against the corresponding gold-standard subset using the distributional metrics above, both per individual fluorescence channel and for the multichannel overlay (composite), to capture channel-specific fidelity as well as cross-channel consistency. StyleGAN2-ADA (62) serves as a GAN-based baseline specialized for limited-data regimes. For each method, 1,000 samples are generated and compared against extracted gold-standard patches. This evaluation is also repeated on both real-world datasets.

##### Missing-channel imputation

One fluorescence channel is withheld and reconstructed from the remaining observed channels; the withheld channel is available as a paired reference and is assessed with the paired metrics (SSIM, LPIPS, Dice). Task-specific baselines are a UNet following the CARE architecture (10, 63), MAXIM (64), and SIMIF (13); together they contrast per-channel and unified imputation designs against the proposed single reliability-conditioned model.

##### Degraded-channel restoration

One channel is degraded at a time, at mild and severe levels, using a physics-based degradation model that combines optical blurring, attenuation, background offset, and Poisson–Gaussian noise (65) (Appendix E). The degraded channel is restored alongside the remaining clean channels, which serve as paired references and are assessed with the same paired metrics. Task-specific baselines are Noise2Void (11) and Cellpose3 (66) (Denoise/Deblur models), while CARE (10) serves as an approximate upper bound. It is trained from scratch on the gold-standard subset using a broad range of degradation parameters spanning both the mild and severe test-time settings, rather than a single fixed configuration. This gives it explicit exposure to the degradation family used at evaluation time. This upper-bound baseline is not used for Col6a1-KI, where the degradation process is unknown and no paired clean reference images are available for training.

##### Joint imputation–restoration

This task combines the two channel-level corruptions above within a single reconstruction: one fluorescence channel is entirely missing while a second is degraded using the same physics-based model (Appendix E, severe degradation level), and both must be recovered jointly from the one remaining reliable channel. To our knowledge, no prior method targets this combined setting, so only the diffusion-based methods introduced above are compared, without task-specific baselines. Two complementary settings are evaluated on BBBC021v1, conditioned on the reliable DAPI channel: degraded Actin together with missing Tubulin, and, with the two roles swapped, degraded Tubulin together with missing Actin. As in the two preceding tasks, the missing and degraded channels are each available as paired references and are assessed with the paired metrics above (SSIM, LPIPS, DICE).

#### Real-world evaluation

On the two rare-disease datasets no paired clean references are available, so both tasks are assessed against surrogate criteria.

##### Dnm1L-Mito (imputation)

Missing-channel imputation is evaluated on the incomplete/non-gold samples: imputed tubulin is compared against the gold-standard tubulin subset using channel-correlation shift (|Δ*ρ*|), tubulin realism statistics (Realism *z*), a Frangi-filter-based (67) fibril-likeness score (Vesselness *z*), and a skeletonized network-topology score (Structure *z*) (Appendix C).

##### Col6a1-KI (restoration)

Degraded-channel restoration is evaluated on tiles for which the Collagen VI channel falls outside *D*_*gold*_, where no paired clean reference exists for the restored channel. Structural fidelity is instead assessed using four complementary scores relative to a gold-standard (real, non-degraded) reference subset: mean vesselness (a Frangifilter-based (67) fibril-likeness score), branch-point count (a skeletonized network-topology descriptor), a topology deviation score (Structure *z*), and the spatial correlation shift between the restored Collagen VI channel and the clean, untouched DAPI channel (Δ*ρ*_DAPI-Col6_), measured relative to its degraded input (Appendix C).

### 4.3. Implementation Details. Model and training

The diffusion backbone is a U-Net (63) with six resolution blocks and two residual layers per block, with a single self-attention block. The diffusion process is defined over *T* = 1000 timesteps using a cosine variance schedule (68), and the auxiliary reconstruction-loss weight is set to *λ*_*x*0_ = 1.0. Reliability masks are drawn per training step from the uniform mixture of reconstruction cases described in Section 3.2. Each fluorescence channel is normalized using global percentile bounds computed on the gold-standard subset, taken as the median across that subset of the per-image 1st and 99th intensity percentiles, and the result is rescaled to [ −1, 1]. Random 256 × 256 crops are extracted from each image and augmented with random scale transformations, except for Dnm1L-Mito, where images are instead resized to 256× 256. Optimization uses AdamW (69) with a base learning rate of 1 × 10^*−*4^ and a cosine schedule with warmup, for 80K iterations at batch size 8. An exponential moving average of the weights with decay 0.999 is maintained during training and used for all reported results, and the final checkpoint is used without validation-based model selection. A single model is trained per dataset on its gold-standard subset *D*_*gold*_, and the same model serves every reconstruction task and inference regime without task-specific retraining. The model is implemented in PyTorch (70) and trained on an NVIDIA RTX 5000 Ada Generation GPU, with an average training time of approximately 4 hours per experiment.

#### Inference

Each inference regime is specified by the triplet (*M, x*_ctx_, *t*_start_) defined in Section 3.2. For degraded-channel restoration, the starting timestep *τ* was fixed by trial rather than swept in a formal ablation: *τ* = *T/*2 = 500 for Col6a1-KI and for the joint imputation–restoration task; for BBBC021v1, *τ* = 200 (mild) / *τ* = 250 (severe) for DAPI restoration, and *τ* = 400 (mild) / *τ* = 500 (severe) for Tubulin and Actin restoration. Resampling/consistency and initial-noise injection were enabled throughout.

#### Baselines

For the comparison methods we use the publicly available implementations whenever available and retain their default architectures and training procedures unless otherwise stated. All baselines are trained on the same gold-standard subset and for the same number of iterations as the proposed model, so that every method sees identical training data. StyleGAN2-ADA (62) is trained using the publicly available implementation, with adaptive discriminator augmentation enabled. For missing-channel imputation, the supervised UNet baseline adopts the CARE architecture (10), trained from scratch with one model per target channel. Since no official SIMIF implementation was available, we reproduced the method from the published description. The baseline DDPM and DDIM variants use the same backbone and training configuration as above but without reliability-mask conditioning; at inference their samplers are adapted to channel imputation by preserving the observed channels throughout the reverse process while sampling only the withheld channel.

## 5. Results

### 5.1. Unconditional Image Generation

We first examine unconditional multichannel generation across BBBC021v1 and the real-world datasets, following the protocol in Section 4.2. Table 2 and Figure 4 present the quantitative and qualitative results, respectively.

**Table 2.** Unconditional generation performance across datasets. Reported per fluorescence channel and for the multichannel overlay using FID (*↓*), KID (*↓*), and iF1 (*↑*), computed on biological embeddings (Appendix C; channel order as in Table 1). For BBBC021v1, the gold-standard reference subset is restricted to 10% of samples (Section 4.1).

| Dataset | Method | Channel 1 |  |  | Channel 2 |  |  | Channel 3 |  |  | Overlay |  |  |
| --- | --- | --- | --- | --- | --- | --- | --- | --- | --- | --- | --- | --- | --- |
| | | FID ↓ | KID ↓ | iF1 ↑ | FID ↓ | KID ↓ | iF1 ↑ | FID ↓ | KID ↓ | iF1 ↑ | FID ↓ | KID ↓ | iF1 ↑ |
| BBBC021v1 | StyleGAN2-ADA | 13.361 | 2.132 | 0.681 | 4.165 | 2.994 | 0.747 | 3.142 | 1.721 | 0.720 | 13.615 | **2.159** | 0.771 |
|  | Baseline DDIM | 8.823 | 1.953 | 0.742 | 7.338 | 3.503 | 0.495 | 7.846 | 3.828 | 0.617 | 21.720 | 7.205 | 0.437 |
|  | Baseline DDPM | 10.655 | **0.826** | 0.881 | 5.001 | 2.478 | **0.940** | 3.989 | 2.478 | **0.930** | 14.042 | 2.504 | **0.900** |
|  | RM-DDIM | 11.625 | 2.684 | 0.624 | 9.522 | 4.481 | 0.467 | 9.202 | 4.640 | 0.441 | 23.895 | 7.855 | 0.499 |
|  | RM-DDPM | **5.850** | 1.240 | **0.925** | **2.420** | **0.880** | 0.899 | **2.271** | **0.940** | 0.918 | **9.454** | 2.631 | 0.522 |
| Col6a1-KI | StyleGAN2-ADA | **15.471** | 9.595 | 0.960 | 9.661 | 4.126 | 0.927 | 6.852 | **1.181** | 0.910 | **15.268** | 7.458 | 0.784 |
|  | Baseline DDIM | 17.522 | 7.088 | 0.513 | **8.317** | **2.403** | 0.925 | **6.444** | 2.409 | 0.907 | 21.176 | 7.290 | 0.110 |
|  | Baseline DDPM | 19.148 | 5.480 | 0.434 | 9.409 | 2.647 | **0.954** | 10.011 | 3.357 | 0.794 | 18.392 | **6.505** | 0.679 |
|  | RM-DDIM | 22.111 | 10.184 | 0.691 | 21.990 | 11.711 | 0.577 | 22.922 | 14.080 | 0.449 | 21.348 | 8.743 | 0.683 |
|  | RM-DDPM | 18.091 | **2.711** | **0.962** | 16.320 | 9.553 | 0.844 | 10.991 | 7.134 | **0.948** | 17.561 | 7.050 | **0.884** |
| Dnm1L-Mito | StyleGAN2-ADA | 10.181 | **1.524** | 0.773 | **4.857** | 2.997 | 0.792 | **7.191** | **2.694** | 0.501 | 9.411 | 2.387 | 0.570 |
|  | Baseline DDIM | **8.974** | 5.635 | 0.632 | 9.698 | **1.463** | **0.965** | 9.776 | 4.213 | 0.560 | **4.901** | **0.832** | 0.875 |
|  | Baseline DDPM | 9.739 | 6.291 | 0.815 | 11.479 | 1.632 | 0.892 | 7.541 | 3.471 | **0.957** | 5.844 | 1.107 | **0.911** |
|  | RM-DDIM | 9.970 | 6.071 | 0.505 | 12.594 | 1.972 | 0.887 | 11.567 | 4.776 | 0.367 | 6.798 | 1.370 | 0.859 |
|  | RM-DDPM | 11.592 | 1.671 | **0.905** | 10.331 | 6.567 | 0.918 | 10.205 | 4.499 | 0.748 | 6.390 | 1.285 | 0.891 |

**Fig. 4.**
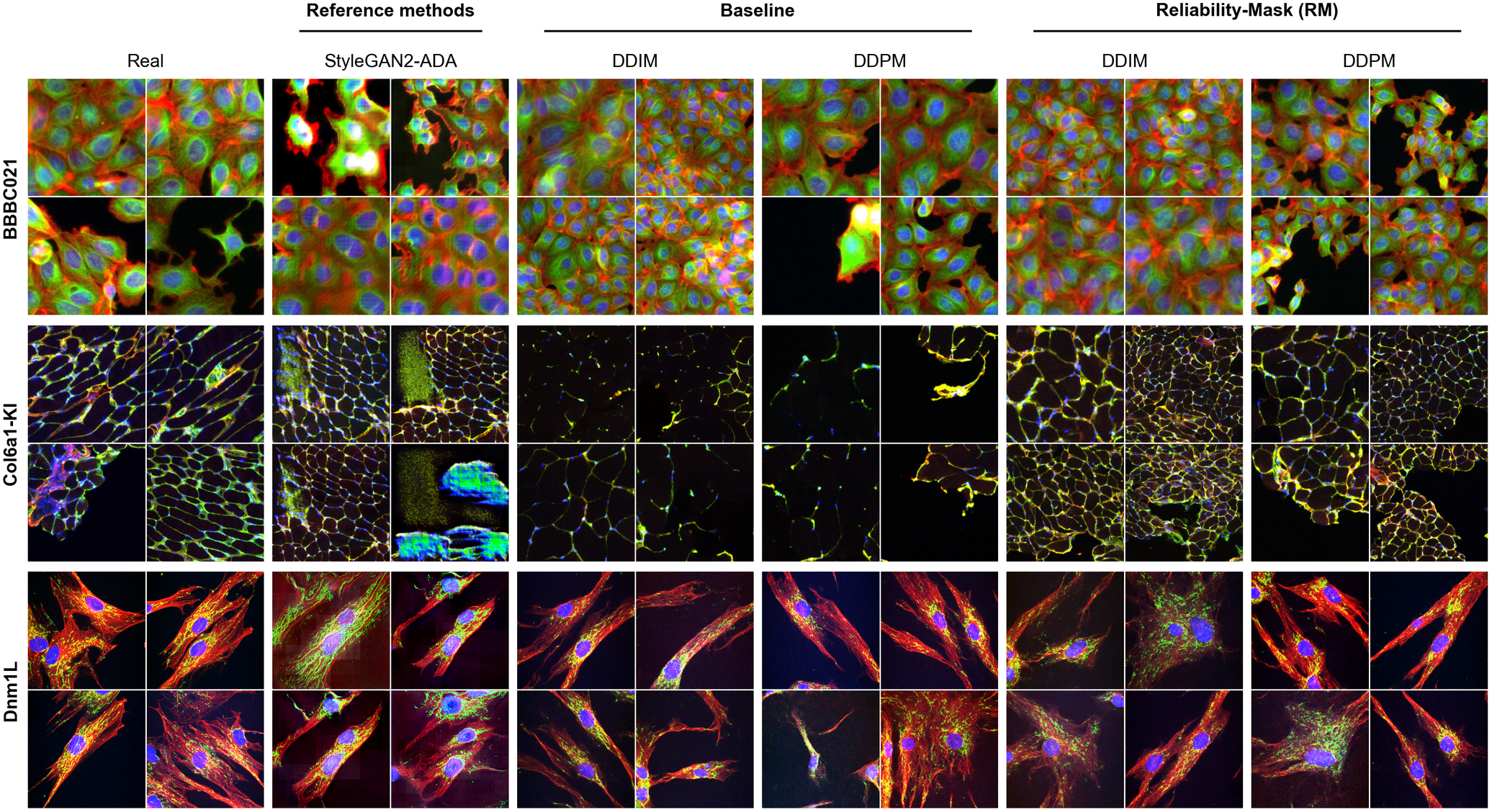
Qualitative unconditional generation results. Real and generated samples for BBBC021v1, Col6a1-KI, and Dnm1L-Mito (rows), compared across StyleGAN2-ADA, Baseline DDIM/DDPM, and RM-DDIM/RM-DDPM (columns). Four samples are shown per method.

RM-DDPM achieves the best FID across all individual channels and the multichannel overlay on BBBC021v1, with per-channel iF1 comparable to Baseline DDPM; its overlay iF1 is nonetheless lower than both Baseline DDPM and StyleGAN2-ADA, echoed by RM-DDIM and Baseline DDIM, suggesting that per-channel fidelity does not guarantee cross-channel realism. On Col6a1-KI, no baseline consistently dominates, with best scores split across StyleGAN2-ADA, Baseline DDIM, and RM-DDPM; only RM-DDIM is consistently worst, recording the highest FID on every channel and the overlay. Qualitative inspection (Figure 4) further shows that StyleGAN2-ADA’s favorable scores coincide with visible checkerboard artifacts, underscoring the limits of distributional metrics alone on this dataset. Results on Dnm1L-Mito are similarly mixed, with no systematic winner across channels and the overlay — consistent with both real-world datasets’ smaller, less curated gold-standard subsets limiting how discriminative these metrics can be.

Qualitative inspection (Figure 4) shows that the diffusion-based methods produce coherent fluorescence structures with fewer visible artifacts across all three datasets, whereas StyleGAN2-ADA exhibits noticeable checkerboard artifacts and local texture inconsistencies, most prominently on BBBC021v1 and Col6a1-KI.

### 5.2. Missing-channel Imputation

We next evaluate missing-channel imputation, where each method reconstructs a held-out fluorescence channel from the remaining input channels, following the protocol in Section 4.2. Tables 3 and 4 report quantitative results on BBBC021v1 and Dnm1L-Mito, respectively, while Figure 5 presents qualitative reconstructions for both datasets.

**Table 3.** Missing-channel imputation performance on BBBC021v1. Mean SSIM (*↑*), LPIPS (*↓*), and DICE (*↑*) per imputed fluorescence channel, over the evaluation set. No. Models indicates whether a method requires a separate model per target channel (3) or a single model for all channels (1).

| Method | No. Models | DAPI |  |  | Tubulin |  |  | Actin |  |  |
| --- | --- | --- | --- | --- | --- | --- | --- | --- | --- | --- |
| | | SSIM ↑ | LPIPS ↓ | DICE ↑ | SSIM ↑ | LPIPS ↓ | DICE ↑ | SSIM ↑ | LPIPS ↓ | DICE ↑ |
| UNet | 3 | 0.729 | 0.437 | 0.859 | **0.528** | 0.573 | 0.820 | 0.543 | 0.536 | 0.824 |
| MAXIM | 3 | 0.758 | 0.372 | 0.864 | **0.528** | 0.560 | 0.830 | **0.554** | 0.510 | 0.826 |
| SIMIF | 1 | 0.353 | 0.502 | 0.747 | 0.170 | 0.566 | 0.731 | 0.195 | 0.567 | 0.783 |
| Baseline DDIM | 1 | 0.485 | 0.448 | 0.639 | 0.393 | 0.483 | 0.770 | 0.409 | 0.491 | 0.759 |
| Baseline DDPM | 1 | 0.471 | 0.440 | 0.648 | 0.354 | 0.482 | 0.767 | 0.353 | 0.500 | 0.756 |
| RM-DDIM | 1 | 0.715 | 0.260 | 0.853 | 0.424 | 0.441 | 0.810 | 0.449 | 0.452 | 0.805 |
| RM-DDPM | 1 | 0.690 | **0.251** | 0.848 | 0.403 | **0.429** | 0.819 | 0.421 | **0.440** | 0.751 |
| RM-DDIM $\times 10$ | 1 | **0.759** | 0.351 | 0.860 | 0.518 | 0.508 | 0.831 | 0.540 | 0.465 | **0.843** |
| RM-DDPM $\times 10$ | 1 | 0.756 | 0.325 | **0.872** | 0.516 | 0.476 | **0.832** | 0.538 | 0.448 | 0.836 |

**Table 4.** Missing-channel imputation performance on Dnm1L-Mito (*α*-tubulin). Surrogate metrics against the gold-standard reference distribution (Section 4.2); no paired ground truth exists for this channel.

| Method | Vesselness $z \downarrow$ | Structure $z \downarrow$ | Realism $z \downarrow$ | $\vert \Delta\rho\vert \downarrow$ |
| --- | --- | --- | --- | --- |
| UNet | 0.939 | 8.411 | 1.188 | 0.050 |
| MAXIM | 0.681 | 4.635 | 1.045 | 0.042 |
| SIMIF | 1.368 | **0.385** | 1.279 | **0.007** |
| Baseline DDIM | 1.731 | 2.493 | 0.926 | 0.247 |
| Baseline DDPM | 1.462 | 2.313 | 0.839 | 0.134 |
| RM-DDIM | 1.109 | 1.644 | 0.801 | 0.084 |
| RM-DDPM | 1.368 | 1.325 | **0.737** | 0.063 |
| RM-DDIM $\times 10$ | **0.563** | 0.593 | 0.982 | 0.081 |
| RM-DDPM $\times 10$ | 0.890 | 0.707 | 0.740 | 0.077 |

**Fig. 5.**
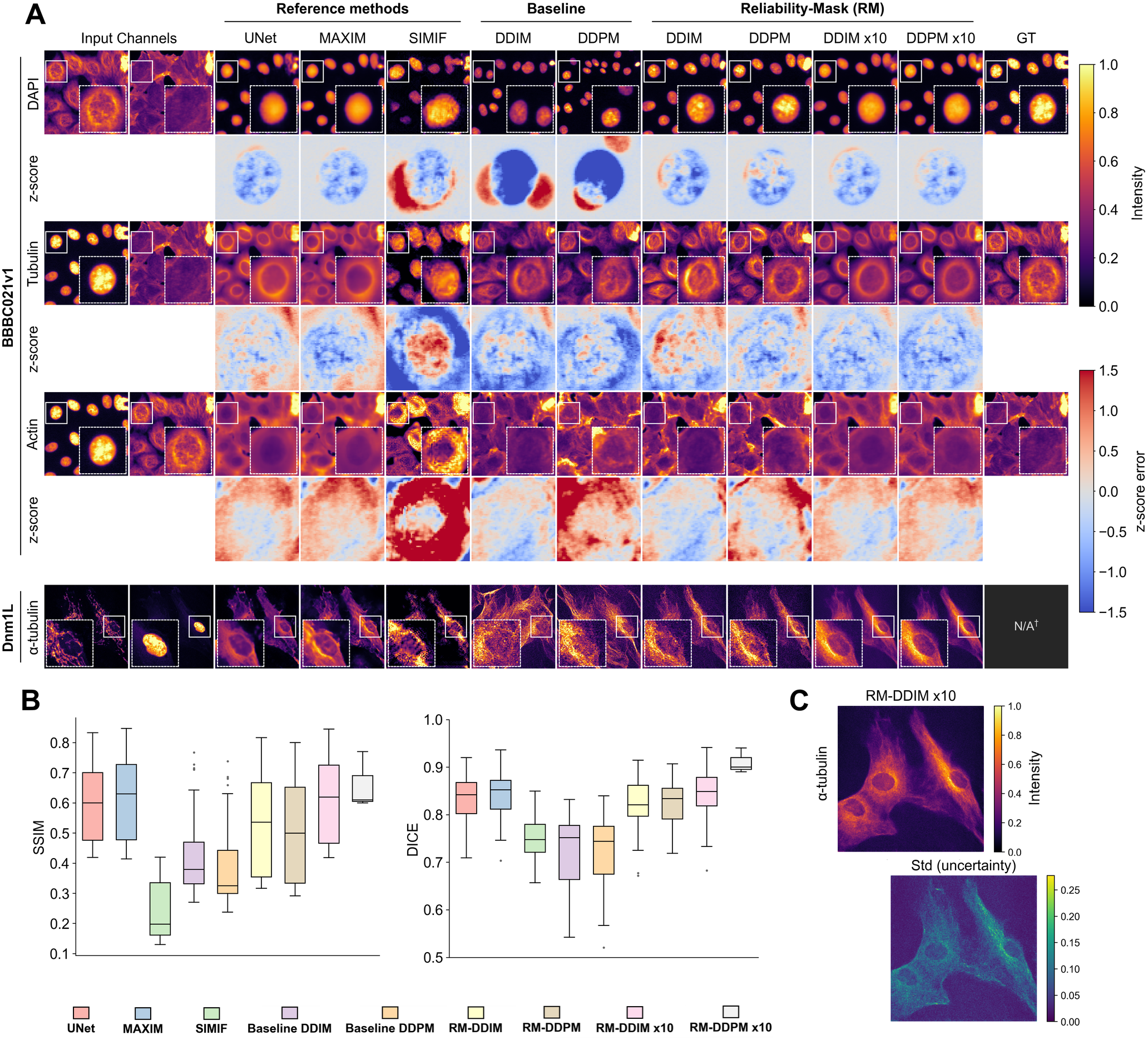
Qualitative missing-channel imputation resultss. (A) Representative reconstructions on BBBC021v1 (DAPI, Tubulin, Actin, each with a z-score error map) and Dnm1L-Mito (*α*-tubulin), across reference (UNet, MAXIM, SIMIF), baseline, and RM methods, including the *×*10 ensemble variants. *†* No paired ground truth is available for this channel. (B) SSIM and DICE distributions across methods. (C) RM-DDIM *×*10 reconstruction and pixel-wise uncertainty (Std) for *α*-tubulin.

On BBBC021v1, no single method dominates across all channels and metrics. Reliability-mask conditioning yields a large gain over the unconditioned baselines: RM-DDIM and RM-DDPM improve SSIM over their Baseline DDIM/DDPM counterparts on every channel (e.g., DAPI: 0.485 →0.715 and 0.471 →0.690), consistent with reliability-mask conditioning mitigating hallucination when reconstructing from the learned prior. RM-DDPM also achieve the best LPIPS consistently across all three channels, indicating superior perceptual fidelity relative to UNet, MAXIM, and SIMIF. SSIM performance is more evenly split: UNet and MAXIM tie for the best Tubulin SSIM, MAXIM achieves the best Actin SSIM, and RM-DDIM ×10 achieves the best DAPI SSIM. DICE, by contrast, is consistently best for the ×10 ensemble variants across all three channels, with RM-DDPM ×10 achieving the best DAPI and Tubulin scores and RM-DDIM ×10 achieving the best Actin score. SIMIF performs worst almost uniformly, with SSIM as low as 0.353 (DAPI) and 0.170 (Tubulin), and LPIPS reaching 0.567 (Actin).

On Dnm1L-Mito, SIMIF yields the best |Δ*ρ*| (0.007) and the best Structure *z* (0.385), but this comes at the cost of the worst Realism *z* (1.279) among all methods, reinforcing the correlation-versus-realism trade-off already visible on BBBC021v1. RM-DDPM achieves the best Realism *z* (0.737), with its ×10 variant close behind (0.740), while also ranking fourth-best on |Δ*ρ*| (0.063), behind only SIMIF, MAXIM, and UNet. Ensembling instead brings the clearest gains on the structural descriptors: RM-DDIM ×10 attains the best Vesselness *z* (0.563) and the second-best Structure *z* (0.593) overall. This benefit does not carry over to Realism *z* symmetrically, however: RM-DDIM degrades substantially under ×10 averaging (0.801 to 0.982), mirroring the LPIPS penalty seen for the ensemble variants on BBBC021v1 above, whereas RM-DDPM’s Realism *z* remains essentially unchanged (0.737 to 0.740). The pixelwise uncertainty map accompanying the RM-DDIM ×10 reconstruction in Figure 5C shows elevated variance concentrated along fine filamentous *α*-tubulin structures, the same regions most affected by the realism/structure trade-off discussed above, indicating that this uncertainty estimate is informative about where the imputed channel is least reliable.

### 5.3. Degraded-channel Restoration

We next evaluate degraded-channel restoration, where each method reconstructs a corrupted fluorescence channel from its degraded observation together with the remaining clean conditioning channels, following the protocol in Section 4.2. Table 5 and Figure 6 present the quantitative and qualitative results on BBBC021v1 under mild and severe degradation.

**Table 5.**
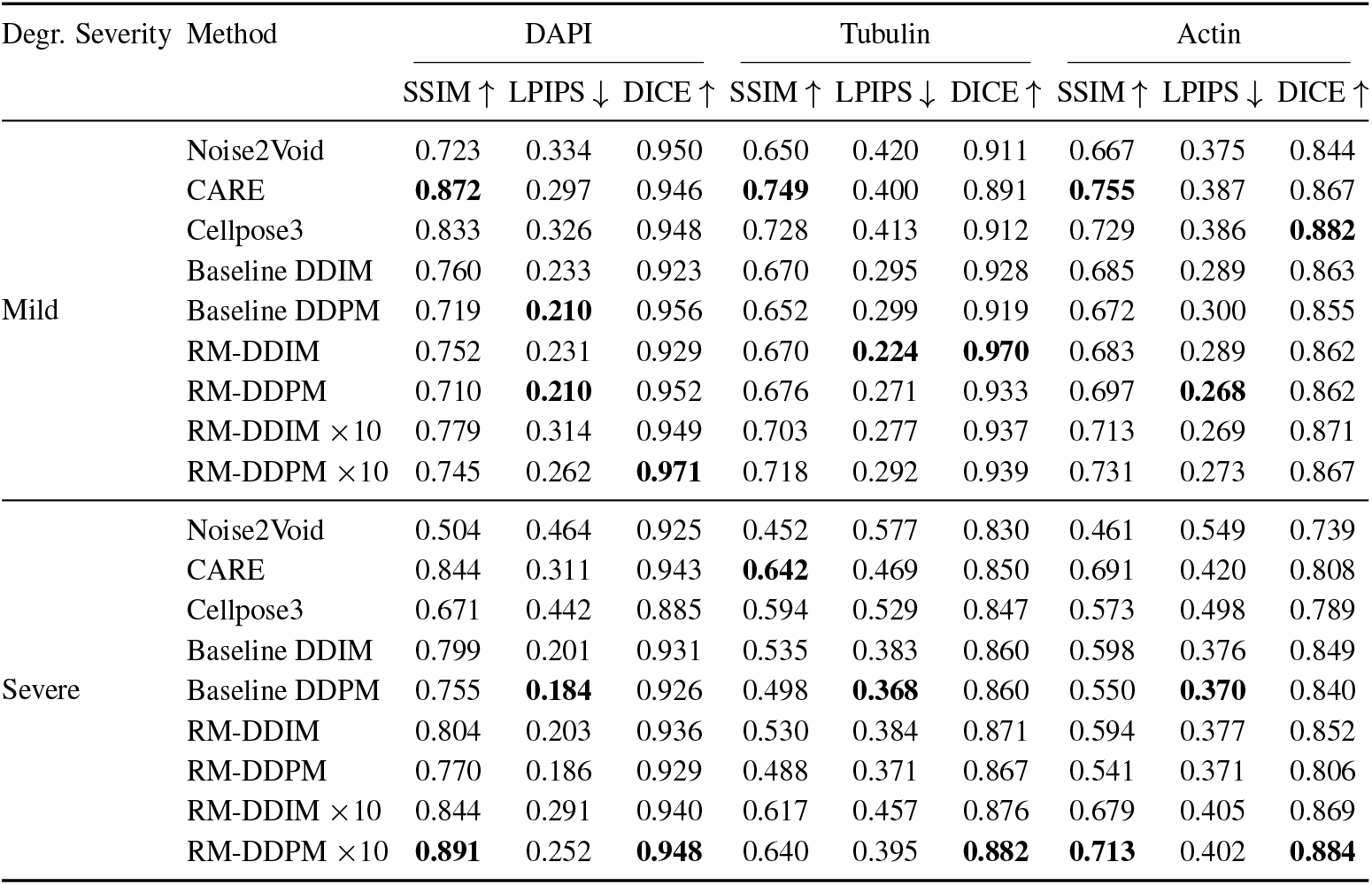
Degraded-channel restoration performance on BBBC021v1. Mean SSIM (*↑*), LPIPS (*↓*), and DICE (*↑*) per fluorescence channel, over the evaluation set. Mild and severe degradation levels follow the physics-based model in Appendix E. Results use the 10% gold-standard subset (Section 4.1).

**Fig. 6.**
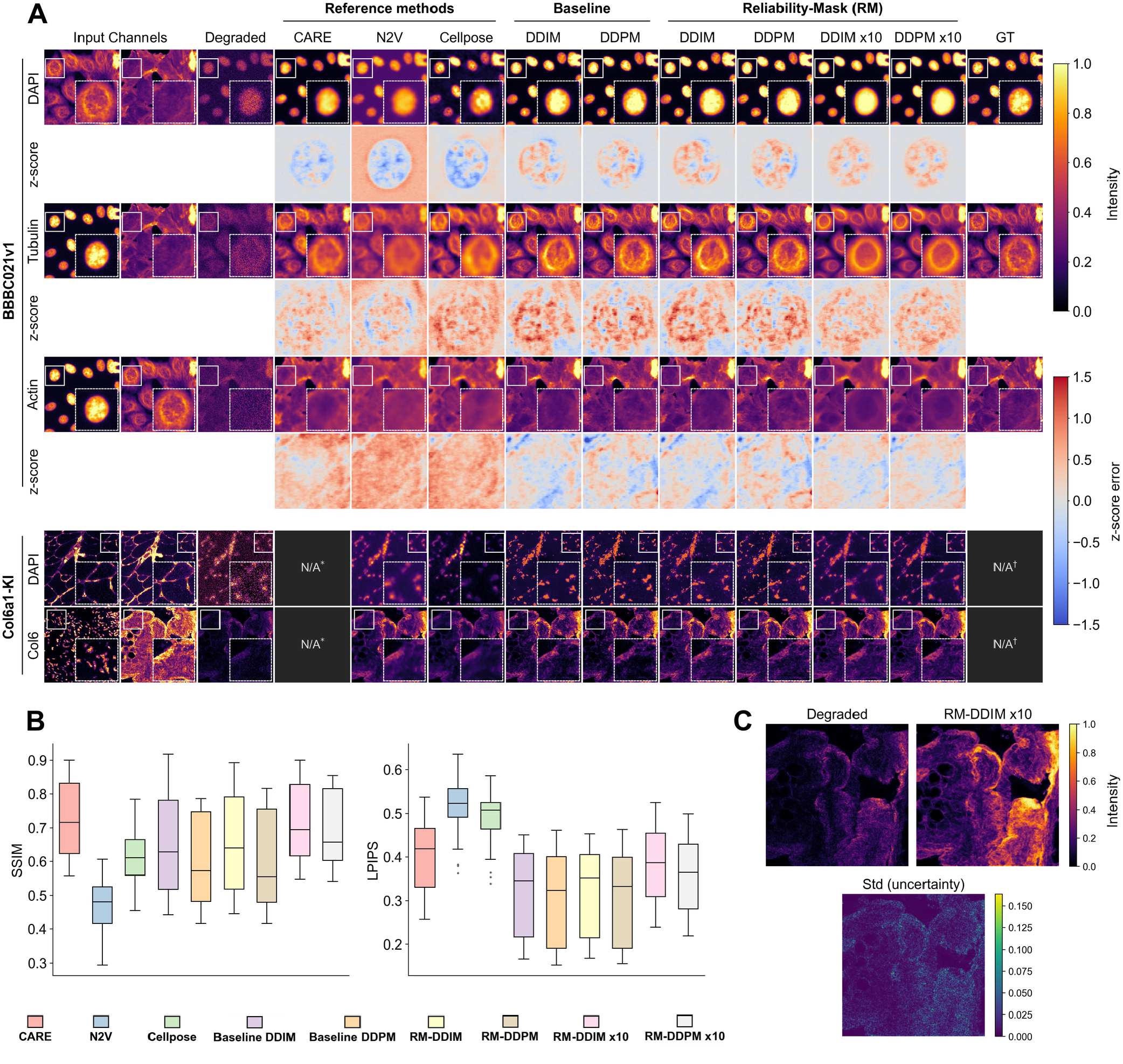
Qualitative degraded-channel restoration results. (A) Representative restorations under severe degradation on BBBC021v1 (DAPI, Tubulin, Actin, each with a z-score error map) and on Col6a1-KI for the DAPI and Collagen VI target channels (quantitative results in Appendix D and Table 6, respectively), across reference, baseline, and RM methods, including the *×*10 ensemble variants. *Method not applicable to this channel. *†* No ground truth is available. (B) SSIM and LPIPS distributions across methods. (C) Degraded input, RM-DDIM *×*10 restoration, and pixel-wise uncertainty (Std) for Collagen VI.

Under mild degradation, CARE achieves the best SSIM across all three channels. Its LPIPS, however, is higher than Baseline DDPM and the single-model RM variants, which instead achieve the best LPIPS on all three channels; DICE shows no single winner, with RM-DDIM best on Tubulin, RM-DDPM ×10 best on DAPI, and Cellpose best on Actin. Under severe degradation, CARE’s SSIM advantage holds only on Tubulin, while RM-DDPM ×10 takes the lead on DAPI and Actin; RM-DDPM ×10 also achieves the best DICE across all three channels, while Baseline DDPM now achieves the best LPIPS across all three channels, reversing the single-model RM advantage seen under mild degradation. As in the previous tasks, the ×10 ensemble variants consistently worsen LPIPS relative to their single-model counterparts under both severity levels, while generally improving SSIM and DICE. Noise2Void performs worst overall, most noticeably under severe degradation.

**Table 6.** Degraded-channel restoration performance on Col6a1-KI (Collagen VI). Surrogate metrics against the gold-standard reference distribution (Section 4.2); no paired ground truth exists for this channel. Gold reference (last row) reports vesselness and branch-point count only; Structure *z* and Δ*ρ* are not applicable.

| Method | Vesselness | Branch pts. | Structure $z \downarrow$ | $\Delta\rho_{\text{DAPI-Col6}} \uparrow$ |
| --- | --- | --- | --- | --- |
| Baseline DDIM | 0.126 | 1525.6 | 0.274 | 0.060 |
| Baseline DDPM | 0.126 | 1540.7 | 0.282 | 0.061 |
| Cellpose3 | 0.072 | 811.1 | 0.387 | 0.030 |
| Noise2Void | 0.099 | 984.8 | 0.352 | 0.032 |
| RM-DDIM | 0.127 | 1646.9 | 0.263 | 0.040 |
| RM-DDPM | **0.127** | **1730.7** | **0.261** | 0.054 |
| RM-DDIM $\times 10$ | 0.115 | 1258.7 | 0.359 | 0.054 |
| RM-DDPM $\times 10$ | 0.115 | 1299.0 | 0.354 | **0.069** |
| Gold reference | 0.173 | 2178.8 | – | – |

Qualitative inspection (Figure 6A) shows that Noise2Void produces the most pronounced z-score error maps, consistent with its lowest median SSIM and highest median LPIPS in Figure 6B, whereas the Baseline DDIM/DDPM and single-model RM-DDIM/RM-DDPM variants show the lowest median LPIPS. The RM-DDIM ×10 and RM-DDPM ×10 variants show the highest median SSIM with the widest spread, mirroring their DICE and SSIM gains in the quantitative results.

On Col6a1-KI, RM-DDPM achieves the closest agreement with the gold reference on both vesselness (0.127 vs. 0.173) and branch-point count (1730.7 vs. 2178.8), and correspondingly the lowest structure *z* (0.261), with RM-DDIM close behind; both single-model RM variants outperform their Baseline DDIM/DDPM counterparts on these topology metrics, mirroring the single-model RM advantage observed on BBBC021v1. The ×10 ensemble variants instead depart further from the gold vesselness and branch-point statistics than their single-model counterparts (e.g., RM-DDPM ×10: 0.115 and 1299.0), while achieving the strongest recovery of the DAPI-Col6 spatial correlation (Δ*ρ*_DAPI-Col6_ = 0.069 for RM-DDPM ×10). This mirrors, rather than reverses, the SSIM/DICE-versus-LPIPS trade-off observed on BBBC021v1, since averaging across samples similarly favors coarse structural consistency (Δ*ρ*) at the expense of fine fibrillar detail (vesselness, branch points). Cellpose3 and Noise2Void perform worst across all four metrics, consistent with their weaker performance on the paired BBBC021v1 restoration task. Qualitatively (Figure 6C), the RM-DDIM× 10 restoration recovers continuous, vessel-like Collagen VI fibrils largely absent from the degraded input, consistent with the elevated Frangi vesselness response in Table 6. The accompanying pixel-wise uncertainty map highlights the recovered fibrils and their immediate surroundings as the least confident regions, consistent with these being the areas most heavily altered relative to the degraded input. This offers a practical signal for flagging restored structures that may warrant closer inspection before downstream quantification.

### 5.4. Joint Channel Imputation and Restoration

We next evaluate the joint setting, where a method must simultaneously impute a missing channel (Tubulin) and restore a degraded channel (Actin), conditioned on the clean DAPI channel, following the protocol in Section 4.2. Table 7 reports quantitative results on BBBC021v1, and Figure 7 presents qualitative reconstructions.

**Table 7.** Joint channel imputation and restoration performance on BBBC021v1. Mean SSIM (*↑*), LPIPS (*↓*), and DICE (*↑*) for the missing (Tubulin) and degraded (Actin) channels, over the evaluation set, conditioned on the reliable

| Method | Tubulin (missing) |  |  | Actin (degraded) |  |  |
| --- | --- | --- | --- | --- | --- | --- |
| | SSIM ↑ | LPIPS ↓ | DICE ↑ | SSIM ↑ | LPIPS ↓ | DICE ↑ |
| Baseline DDIM | 0.320 | 0.540 | 0.712 | 0.417 | 0.491 | 0.722 |
| Baseline DDPM | 0.302 | 0.526 | 0.707 | 0.387 | 0.497 | 0.672 |
| RM-DDIM | 0.397 | 0.484 | 0.747 | 0.422 | 0.476 | 0.580 |
| RM-DDPM | 0.371 | **0.463** | 0.762 | 0.404 | 0.469 | 0.576 |
| RM-DDIM $\times 10$ | **0.496** | 0.523 | 0.764 | **0.516** | **0.456** | 0.712 |
| RM-DDPM $\times 10$ | 0.469 | 0.512 | **0.775** | 0.510 | 0.465 | **0.753** |

**Fig. 7.**
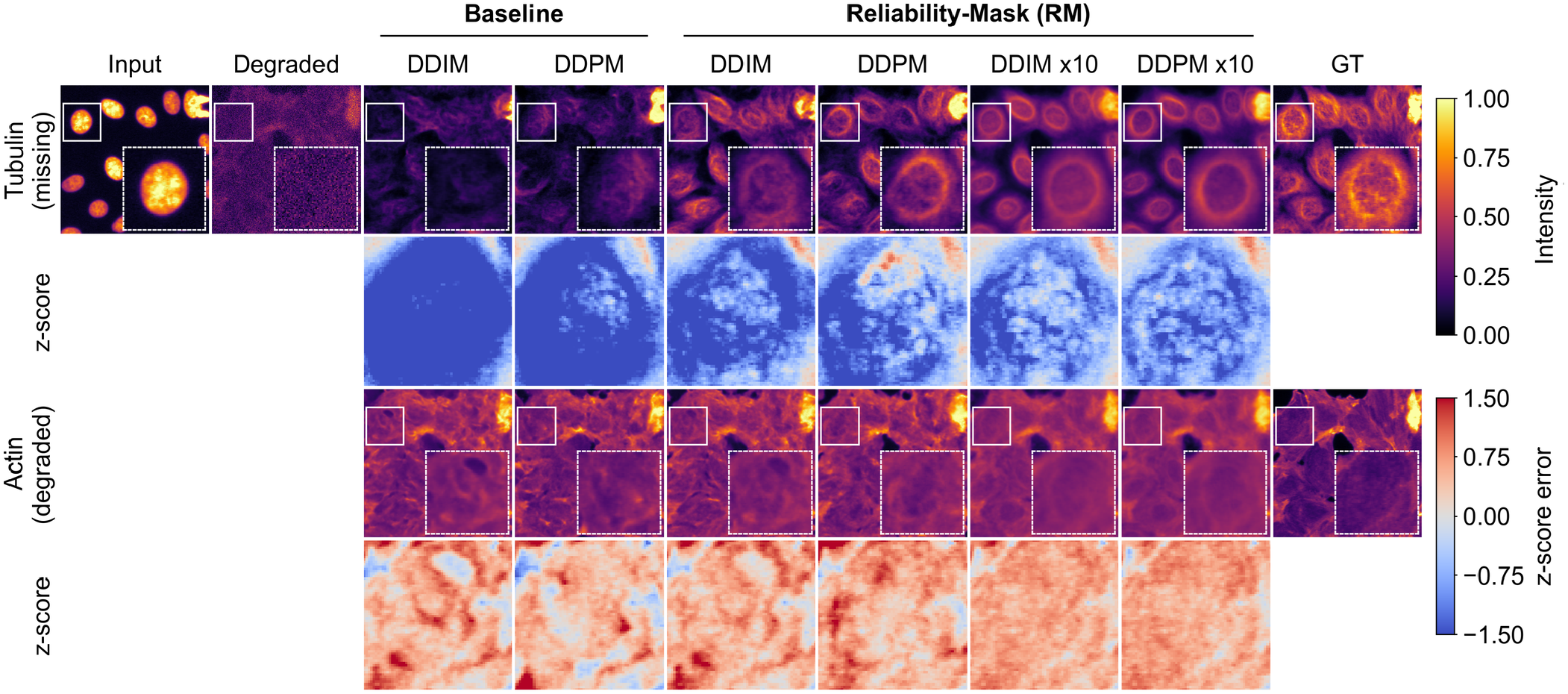
Qualitative joint channel imputation and restoration results. Joint reconstruction of the missing Tubulin and degraded Actin channels, conditioned on the clean DAPI channel, on BBBC021v1, across baseline and RM methods, including the *×*10 ensemble variants; each channel is shown with a z-score error map.

RM-DDIM ×10 achieves the best SSIM on both the missing Tubulin channel (0.496) and the degraded Actin channel (0.516), as well as the best LPIPS on Actin (0.456). RM-DDPM achieves the best LPIPS on Tubulin (0.463), the only metric on which a single-model variant outperforms the ×10 ensemble. Both RM variants outperform their Baseline DDIM/DDPM counterparts on SSIM for both channels, consistent with reliability-mask conditioning mitigating hallucination relative to the unconditioned baselines. RMDDPM × 10 trails RM-DDIM ×10 on this case, but achieves the best SSIM of all compared methods on the complementary case (missing Actin, degraded Tubulin; Appendix F). RM-DDPM ×10 achieves the best DICE on both channels (0.775 on Tubulin, 0.753 on Actin), while the single-model RM variants score below both baselines on Actin (0.576–0.580 vs. 0.672–0.722), mirroring the SSIM/DICE-versus-LPIPS ensembling trade-off observed in the imputation and restoration tasks above.

## 6. Discussion

RM-DDPM formulates multichannel generation, imputation, and restoration as instances of a single reliability-conditioned inference problem (Section 3.2). A single model, driven only by the reliability mask *M* and starting timestep *t*_start_, replaces the per-task and per-channel models required by UNet, MAXIM, and SIMIF (*No. Models* column, Table 3). Despite this reduced specialization, it is competitive with, and on several channels and metrics surpasses, these task-specific baselines (Tables 3–5). The same stochastic sampling also yields pixel-wise uncertainty maps as a by-product (Section 3.4; Figures 5C and 6C). This provides a confidence signal for flagging unreliable reconstructions that none of the compared baselines offer.

The results further indicate that a small, curated reference set can substantially reduce data requirements. RM-DDPM/RM-DDIM remained competitive using only 10% of BBBC021v1 as gold-standard, and 113 tiles from 23 Col6a1-KI mosaics (Section 4.1). This is consistent with rare-disease settings, where large uniformly clean cohorts are rarely available (Section 1). It suggests that an existing dataset’s value depends less on its size than on whether a compact high-quality subset can be identified. The benefit of reliability-mask conditioning tracks how scarce this subset is. On Dnm1L-Mito (95% gold), Baseline DDPM already matches or exceeds RM-DDPM on several metrics (Table 2). The RM variants instead show a clearer advantage on BBBC021v1 and Col6a1-KI (10% and 12.5% gold).

A related trend is that this advantage over supervised baselines grows with degradation severity. CARE leads SSIM under mild degradation, but RM-DDPM × 10 takes over under severe degradation and on DICE overall (Table 5), consistent with a learned prior mattering most when the observed signal is least informative. A similar pattern holds across dataset scale and channel count: baselines relying on cross-channel redundancy and large training sets are most disadvantaged on these small, three-channel datasets, where inter-channel correlation is comparatively weak and few samples are available to learn from (Section 2).

These gains are not unconditional, however, and several limitations qualify them.

### Limitations

Performance is bounded by the gold-standard subset used for training and normalization (Section 4.3). Its quality-control criteria were calibrated from expert-labeled exemplars for Col6a1-KI but applied automatically without further review (Appendix A), while the BBBC021v1 gold subset is a random rather than quality-based sample; neither has been validated against full expert curation. A foundation model pretrained exclusively on large, curated gold-standard multichannel fluorescence datasets would reduce this dependence on small, dataset-specific curation, lower the per-dataset training cost, and likely improve performance, though such large collections remain scarce, particularly for rare-disease imaging.

Pixel-space sampling over *T* = 1000 steps is also costly relative to the single-pass baselines (UNet, MAXIM, Noise2Void, Cellpose3). Hallucination also persists, most visibly in imputation: ×10 ensembling improves SSIM/DICE but worsens LPIPS/realism (Sections 5.2, 5.3), because averaging multiple stochastic samples smooths out high-frequency detail at the expense of perceptual realism. The unconditional restoration regime, most exposed to this risk, is not separately evaluated with quantitative metrics (Section 5).

All metrics used here assess structural, perceptual, or distributional agreement rather than photometric accuracy, so reconstructed intensities should not be treated as quantitatively reliable for downstream intensity-based analysis until such calibration is validated. Distributional generation metrics (FID, KID, iF1) are themselves known to become unreliable with small reference sets (61, 71), such as those available for Col6a1-KI and Dnm1L-Mito, calling for evaluation protocols better suited to small-sample generative comparisons; they are also not always mutually consistent, as StyleGAN2-ADA obtained the best FID/KID on Col6a1-KI despite visibly artifactual samples (Figure 4).

### Future work

Diffusion in a learned embedding space would reduce the computational cost of pixel-space sampling. Pretraining a foundation model exclusively on large, curated gold-standard multichannel fluorescence collections is a promising direction for reducing both curation and per-dataset training costs. Combining the current automated quality-control pipeline with expert-in-the-loop review would help validate the gold-standard subset, and comparing it against random subset selection would further clarify how curation strategy affects performance. Quantitatively evaluating the unconditional-restoration and joint regimes would better characterize hallucination risk where context is least reliable. An intensity-calibration mechanism would extend the framework toward quantitative downstream analysis. Developing evaluation protocols better suited to small-sample generative comparisons, and combining embedding-based scores with human-in-the-loop assessment, would help resolve cases where FID/KID/iF1 are unreliable or disagree with qualitative inspection.

## Data Availability

BBBC021v1 is a publicly available dataset from the Broad Bioimage Benchmark Collection (51, 52), freely accessible at https://bbbc.broadinstitute.org/BBBC021. The Col6a1-KI and Dnm1L-Mito datasets contain fluorescence microscopy images derived from patient-related biological material collected at Hospital Sant Joan de Déu and cannot be made publicly available.

## Code Availability

Code implementing RM-DDPM, the baselines, and the evaluation protocol used in this work is publicly available at https://github.com/igeor/ddpm4het.

## ACKNOWLEDGEMENTS

This work was funded in part by the European Union under HORIZON-MSCA-2022-DN project: Improving BiomEdical diagnosis through LIGHT-based technologies and machine learning “BE-LIGHT” (GA no. 101119924 – BE-LIGHT).

## Appendix A: Gold-Standard Subset Selection and Quality Assessment

For the Col6a1-KI dataset, the channel-specific IQMs and thresholds used in Φ_*c*_ were derived from an expert-guided calibration procedure. For each fluorescence channel, experts selected representative examples with clearly high-quality and clearly low-quality signal. These examples were used only for quality-control calibration and were not used for diffusion-model training.

A broad set of candidate IQMs was computed for the expert-selected images. The candidate set included mean intensity, standard deviation, low and high intensity percentiles, percentile spread, interquartile range, coefficient of variation, robust coefficient of variation, skewness, kurtosis, signal-to-background ratio, signal-to-noise proxy, saturation rate, low-signal rate, zero fraction, maximum-value fraction, dynamic-range usage, Laplacian-variance sharpness, Tenengrad sharpness, and Brenner focus.

Threshold discovery was performed independently for each channel. For each channel, the expert-labelled high-quality and low-quality examples were treated as two classes, and a shallow decision tree was fitted using the candidate IQMs as input features. The tree was used as an interpretable threshold-selection tool to identify the subset of IQMs and threshold values that best separated the two expert-defined groups.

The resulting channel-specific criteria were then used as the predefined quality-control rules for the Col6a1-KI dataset. The selected IQMs and thresholds are reported in Table 8.

**Table 8.** Channel-specific IQMs and thresholds used for Col6a1-KI gold-standard selection. A sample was retained only if all criteria were satisfied for all required channels.

| Channel | IQM | Criterion |
| --- | --- | --- |
| DAPI | Laplacian-variance sharpness | $< 0.10$ |
| DAPI | Skewness | $> 4.0$ |
| DAPI | Coefficient of variation | $> 2.1$ |
| DAPI | Interquartile range | $< 0.04$ |
| Perlecan | Standard deviation | $> 0.10$ |
| Perlecan | Percentile spread | $> 0.5$ |
| Perlecan | Tenengrad sharpness | $> 0.011$ |
| Perlecan | Brenner focus | $> 0.007$ |
| Collagen VI | Standard deviation | $> 0.125$ |
| Collagen VI | Percentile spread | $> 0.6$ |

The selected criteria also indicate the main degradation modes present in the Col6a1-KI data. For DAPI, they mainly reject images with unstable background, loss of clear nuclear sparsity, or high-frequency artifact-like texture. For Perlecan, they reject low-contrast and poorly focused images. For Collagen VI, they mainly reject weak-signal or low-dynamic-range images. Overall, the QC rules target residual degradation related to weak fluorescence signal, reduced contrast, blur, and unstable background.

## Appendix B: Reliability Mask Sampling

The masking distribution is defined over seven non-trivial reconstruction cases, obtained by excluding the identity case *M* = **1**:

1. small spatial mask in one channel,
2. small spatial mask in multiple channels,
3. large spatial mask in one channel,
4. large spatial mask in multiple channels,
5. full masking of one channel,
6. full masking of multiple channels,
7. full masking of all channels.

At each optimization step a single case is drawn uniformly at random, so that all cases are sampled with equal probability throughout training:

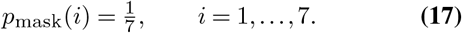

This uniform mixture exposes the model to channel-level missingness, localized spatial unreliability, and their combinations in equal proportion, without favouring any particular reconstruction task. Case 7 (*M* = **0**) is retained so that the model also learns the unconditional multichannel prior.

Spatial masks are sampled as rectangular regions. Small masks cover a random area fraction in [0.20, 0.35], while large masks cover a random area fraction in [0.50, 0.70]. Rectangle aspect ratios are sampled randomly to avoid degenerate strip-like masks.

## Appendix C: Biological Feature Embeddings and Real-World Surrogate Metrics

### Biological feature embedding space

For each fluorescence channel, a per-channel foreground mask is obtained via Otsu thresholding (72) (after removing small spurious objects), and channel-specific descriptors are computed from it: for the nuclei channel, foreground area, object circularity, mean intensity, texture contrast, and object count; for the actin channel, foreground area, object eccentricity and solidity, mean intensity, and texture contrast; for the tubulin channel, mean intensity, a radial intensity-distribution score, texture contrast, and a granularity score capturing high-frequency structure. Texture contrast is computed from a gray-level co-occurrence matrix (73). A small set of cross-channel descriptors is appended when the relevant channels are present, including area and intensity ratios between channels and the spatial overlap between their foreground masks. Concatenating these descriptors yields a single feature vector per image patch, which serves as the embedding space for all generative-quality metrics reported in the main text (iF1, FID, KID). Patches are 256 × 256 px and are normalized using the same percentile-based intensity normalization applied during training (Section 4.3) prior to descriptor extraction.

Per-channel and multichannel-overlay views share this same descriptor pipeline: the overlay feature vector concatenates all per-channel and cross-channel descriptors, whereas each per-channel view retains only the descriptors for that channel, since the cross-channel terms require multiple simultaneously present channels.

#### Distributional metrics (iF1, FID, KID)

iF1, FID, and KID are computed in this feature space analogously to their standard Inception-embedding counterparts (59–61): real and generated feature sets are jointly standardized, iF1 uses adaptive *k*-nearest-neighbor manifold radii (*k* = 3) to estimate precision and recall, and FID is the Fréchet distance between the fitted Gaussians. KID is an unbiased maximum mean discrepancy with a cubic polynomial kernel,

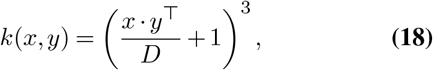

where *D* is the feature dimensionality.

#### Real-world surrogate metrics (Dnm1L-Mito and Col6a1-KI)

For the two rare-disease datasets, where no paired clean reference is available, the following surrogate scores are used instead.

##### *Channel-correlation shift (* |Δ*ρ*|, *Dnm1L-Mito)*. Compares the cross-channel Pearson correlation between the imputed tubulin channel and the other observed channels against the same correlation computed on gold-standard samples, capturing whether imputation preserves realistic cross-channel dependency

##### Realism z

Summarizes the imputed channel’s intensity, contrast, entropy, and foreground coverage into five descriptors *f*, and reports their mean deviation from the gold-standard distribution in standard-score units,

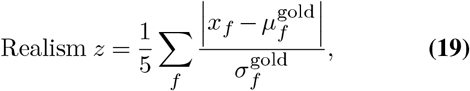

penalizing outputs that are statistically implausible even when cross-channel correlation is preserved.

*Vesselness z*. A multiscale Frangi vesselness filter (67) is applied to the imputed tubulin channel to enhance fibril-like structures, and its mean response over the foreground mask, *v*, is compared against the gold-standard distribution,

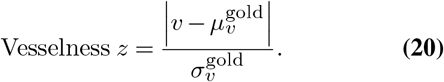

For Col6a1-KI (Table 6), the same mean vesselness response *v* is reported directly as a raw value alongside an explicit gold-reference row, rather than as a z-score, since the gold-standard distribution can be shown in the table itself.

##### Structure z

The foreground mask of the imputed or restored target channel is skeletonized (74), and a small set of network-topology descriptors *g* (branch-point count and total skeleton length) is extracted from the skeleton. Their mean deviation from the gold-standard distribution is reported analogously,

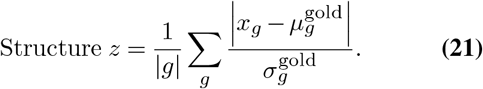

For Col6a1-KI, branch-point count is additionally reported on its own as a raw value against the gold-reference row, alongside Structure *z*.

##### *Correlation shift (*Δ*ρ*_*DAPI-Col6*_, *Col6a1-KI)*. A distinct quantity from the gold-deviation score |Δ*ρ*| used for Dnm1L-Mito above: it is the Pearson correlation between the restored Collagen VI channel and the clean, untouched DAPI channel, measured relative to the same correlation computed on the degraded (pre-restoration) input, capturing whether restoration recovers realistic cross-channel dependency rather than deviation from a gold population

All Dnm1L-Mito real-world surrogate metrics (|Δ*ρ* |, Realism *z*, Vesselness *z*, and Structure *z*) are computed on a reduced *n* = 4 evaluation subset for which skeletonizable, well-segmented tubulin masks could be reliably extracted, and should therefore be interpreted as exploratory rather than definitive.

## Appendix D: DAPI Restoration Quality on Col6a1-KI

As a complement to the Collagen VI restoration results in Section 5.3 (Table 6), Table 9 reports degraded-channel restoration performance on the DAPI channel of Col6a1-KI, following the same evaluation protocol (Section 4.2). Since the DAPI signal on this dataset consists of small, sparse puncta rather than large nuclear disks, structural fidelity is instead assessed with density-based descriptors: mean blob area and nearest-neighbor distance (NND), alongside the topology deviation score (Structure *z*) and the spatial correlation shift between the restored DAPI channel and the clean, untouched Perlecan channel (Δ*ρ*), measured relative to its degraded input.

**Table 9.** Degraded-channel restoration performance on Col6a1-KI (DAPI). Surrogate metrics against the gold-standard reference distribution (Section 4.2); no paired ground truth exists for this channel. Gold reference (last row) reports blob area and NND only; Structure *z* and Δ*ρ* are not applicable.

| Method | Blob area (px) | NND (px) | Structure $z \downarrow$ | $\Delta\rho \uparrow$ |
| --- | --- | --- | --- | --- |
| Baseline DDIM | 845.9 | 20.4 | 0.489 | 0.560 |
| Baseline DDPM | 1038.9 | 25.7 | 0.970 | 0.545 |
| Cellpose3 | **305.1** | 20.6 | **0.243** | 0.538 |
| Noise2Void | 320.6 | 21.8 | 0.341 | 0.543 |
| RM-DDIM | 448.6 | 19.5 | 0.336 | 0.571 |
| RM-DDPM | 464.5 | **18.3** | 0.300 | **0.577** |
| RM-DDIM $\times 10$ | 584.0 | 23.6 | 0.488 | 0.560 |
| RM-DDPM $\times 10$ | 630.6 | 25.3 | 0.635 | 0.561 |
| Gold reference | 79.9 | 17.8 | – | – |

RM-DDPM achieves the closest agreement with the gold reference on both NND (18.3 vs. 17.8) and DAPI–Perlecan spatial correlation (Δ*ρ* = 0.577), with RM-DDIM close behind; both single-model RM variants outperform their Baseline DDIM/DDPM counterparts on these two metrics, mirroring the single-model RM advantage observed on BBBC021v1 and on Collagen VI (Section 5.3). Cellpose3 instead attains the lowest blob area (305.1, closest to the gold 79.9) and the lowest structure *z* (0.243), consistent with its per-object segmentation being well suited to the small, sparse puncta that characterize the DAPI signal on this dataset; this advantage does not carry over to Δ*ρ*, where Cellpose3 scores lowest among all methods. The ×10 ensemble variants depart further from the gold blob-area and structure-*z* statistics than their single-model counterparts and, unlike on Collagen VI, do not recover this loss through improved Δ*ρ* either, trailing their single-model counterparts on all four metrics here.

## Appendix E: Fluorescence Image-Formation Degradation Model

Let *x* ∈ [0, 1]^*H×W*^ denote a single clean fluorescence channel. The degradation operator applies four sequential stages — optical blur, attenuation with background offset, photon shot noise, and detector read noise (65) — parameterized by a severity level *s* ∈{mild, severe}.

### Optical blur

A normalized 2D Gaussian point-spread-function approximation (75) is applied via convolution:

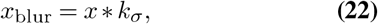

where *k*_*σ*_ is a separable 2D Gaussian kernel of standard deviation *σ* (`blur_sigma`) and odd kernel size ⌊6*σ* + 1⌋.

### Attenuation and background offset

The blurred signal is scaled by a constant attenuation factor *a* and shifted by a constant additive background level *b*:

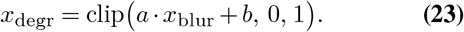

### Photon shot noise

A Poisson noise process is applied at an effective photon-count scale *γ* (`photon_scale`), where lower *γ* corresponds to stronger shot noise:

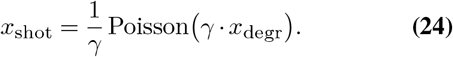

### Detector read noise

Zero-mean additive Gaussian noise with standard deviation *σ*_*r*_ (`read_noise_std`) is added, and the result is clipped to the valid intensity range:

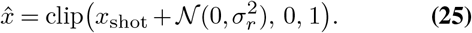

The parameter set (*σ, a, b, γ, σ*_*r*_) is fixed per severity level for the evaluation-time degradation operator, as summarized in Table 10.

**Table 10.**
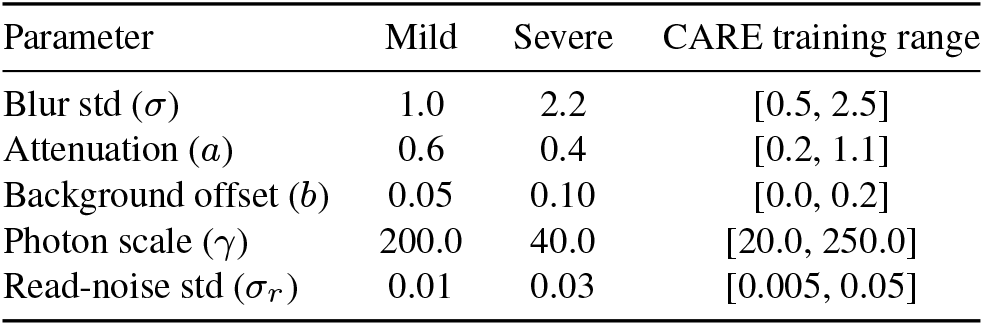
Degradation parameters used at evaluation time for the mild and severe severity levels, together with the ranges from which degradations were sampled when training the CARE upper bound.

| Parameter | Mild | Severe | CARE training range |
| --- | --- | --- | --- |
| Blur std ($\sigma$) | 1.0 | 2.2 | [0.5, 2.5] |
| Attenuation ($a$) | 0.6 | 0.4 | [0.2, 1.1] |
| Background offset ($b$) | 0.05 | 0.10 | [0.0, 0.2] |
| Photon scale ($\gamma$) | 200.0 | 40.0 | [20.0, 250.0] |
| Read-noise std ($\sigma_r$) | 0.01 | 0.03 | [0.005, 0.05] |

## Appendix F: Joint Channel Imputation and Restoration: Complementary Case

As a complementary check on the joint imputation– restoration task of Section 5.4, the two corruption roles are swapped: Actin is entirely missing while Tubulin is synthetically degraded, again conditioned on the clean DAPI channel and evaluated on the same BBBC021v1 test set and paired metrics (SSIM, LPIPS, DICE). Table 11 reports the results. RM-DDPM ×10 achieves the best SSIM of all compared methods on both channels (0.465 on the missing Actin channel, 0.534 on the degraded Tubulin channel), reversing its underperformance relative to RM-DDIM ×10 on the main-text case (Section 5.4). RM-DDIM ×10 retains the best LPIPS on the degraded channel (0.442), while RM-DDPM (single-model) attains the best LPIPS on the missing channel (0.492).

**Table 11.** Joint channel imputation and restoration performance on BBBC021v1, complementary case. Mean SSIM (*↑*), LPIPS (*↓*), and DICE (*↑*) for the missing (Actin) and degraded (Tubulin) channels, over the evaluation set, conditioned on the clean DAPI channel.

| Method | Actin (missing) |  |  | Tubulin (degraded) |  |  |
| --- | --- | --- | --- | --- | --- | --- |
| | SSIM ↑ | LPIPS ↓ | DICE ↑ | SSIM ↑ | LPIPS ↓ | DICE ↑ |
| Baseline DDIM | 0.282 | 0.535 | 0.477 | 0.401 | 0.487 | 0.797 |
| Baseline DDPM | 0.343 | 0.511 | 0.464 | 0.391 | 0.472 | 0.785 |
| RM-DDIM | 0.346 | 0.497 | 0.473 | 0.398 | 0.459 | 0.796 |
| RM-DDPM | 0.384 | **0.492** | 0.533 | 0.406 | 0.455 | 0.785 |
| RM-DDIM $\times 10$ | 0.425 | 0.529 | **0.672** | 0.511 | **0.442** | **0.819** |
| RM-DDPM $\times 10$ | **0.465** | 0.512 | 0.651 | **0.534** | 0.472 | 0.805 |

